# Mechanism of repression of the glycerol utilization operon in *Borrelia burgdorferi*

**DOI:** 10.1101/2022.11.01.514788

**Authors:** Jun-Jie Zhang, Sajith Ranghunandanan, Qian Wang, Yongliang Lou, X. Frank Yang

## Abstract

Lyme disease pathogen *Borrelia burgdorferi*, switches carbohydrate utilization while transmitting between two disparate hosts, an *Ixodes* tick vector and a mammalian host. The ability to use glycerol is important for *B. burgdorferi* to colonize and persist in the tick vector, and expression of the *glpFKD* (*glp*) operon for glycerol uptake/utilization is tightly regulated during the enzootic cycle of *B. burgdorferi* between ticks and mammals. In present study, we identified a *cis* element within the 5’ untranslated region of *glp* that negatively regulates the *glp* expression. This repression of *glp* is independent of RpoS. We further show that BadR directly binds to this *cis* element and represses *glp*. The efficiency of BadR binding in the presence of c-di-GMP and various carbohydrates were also assessed. This finding, along with previous findings of positive regulation of *glp* expression by the c-di-GMP signaling pathway and negative regulation by the alternative sigma factor RpoS, demonstrates that *B. burgdorferi* employs multi-layer regulatory mechanisms to coordinate *glp* expression during its enzootic cycle.

## INTRODUCTION

*Borrelia* burgdorferi, a spirochetal pathogen that causes Lyme disease, is maintained in an enzootic cycle involving an arthropod vector, *Ixodes* ticks, and a mammalian reservoir host, such as field mice, squirrels, and birds (1, 2). Despite its diverse host range and complicated life cycle, *B. burgdorferi* as an obligated pathogen has a reduced genome with a very restricted metabolic capacity (3-6). As for most bacteria, glucose is the preferred carbohydrate for *B. burgdorferi*, and it is highly available during its mammalian phase of the cycle. In the tick phase of the cycle, *B. burgdorferi* is confined to the nutrient-poor midgut lumen of the unfed nymph for months (7-11). Glycerol is produced by certain insects as well as arthropods as a cryoprotective molecule (12, 13). The ability of *B. burgdorferi* to utilize glycerol as a carbohydrate source is critical for spirochete’s maximal fitness in ticks (4, 9, 13, 14).

The *glpFKD* (*glp*) operon of *B. burgdorferi* encodes GlpF (Bb0240), a transmembrane facilitator protein for the uptake of glycerol into the cell; GlpK (Bb0241), a kinase that phosphorylates glycerol to glycerol 3-phosphate (G3P); and GlpD (Bb0243), a G3P dehydrogenase that converts G3P to dihydroxyacetone phosphate (4). Spirochetes with *glp* deficiency significantly reduced survival in ticks and lowered transmission rate to mammals via tick bite (9, 11, 13). *B. burgdorferi* tightly regulates its *glp* expression during the tick-mammal cycle in response to nutritional and environmental factors such as carbon source (i.e., glycerol) and temperature (9, 11,13, 15, 16). Previously we and others showed that *glp* expression is under the control of the Hk1-Rrp1 two-component signaling system, a system that produces the second messenger cyclic di-GMP. The Hk1-Rrp1 system is exclusively required for spirochetes to survive in the tick phase of the enzootic cycle (9, 17, 18). A *rrp1* mutant or a mutant lacking the c-di-GMP effector PlzA has defect in *glp* expression level and has low growth rate in glycerol as a sole carbon source(9, 17) (11, 19). In addition to c-di-GMP, other factors such as (p)ppGpp also showed regulating *glp* expression. (4, 9, 11, 14, 16, 19-23).

When *B. burgdorferi* transmits from ticks to mammals, it activates genes required for mammalian infection while repressing genes for tick adaptation. The alternative sigma factor RpoS plays a central role in this reciprocal regulation of mammalian-phase and tick-phase gene expression of *B. burgdorferi*. The *glp* expression is repressed under mammalian host-adapted conditions and such repression depends on RpoS (13). Grove et al. recently showed that RpoS directly represses *glp* expression by competing with the housekeeping σ^70^ promoter (35). In this study, we identified a *cis*- and a *trans*-element that represses *glp* expression independent of RpoS.

## RESULTS

### Identification of the *cis* element involved in the negative regulation of *glp* expression

Previous reports show that the transcriptional start site of the *glp* is located at 195 bp upstream from the ATG codon, resulting in a long untranslated leader sequence (UTR) (24, 25). Bioinformatics analysis (BPROM, http://linux1.softberry.com/berry.phtml) revealed the presence of another -35/-10 promoter sequence located within the leader sequence (**Fig. 1A**). To investigate the function of these two -35/-10 promoter sequences, three shuttle vectors were constructed: one carrying both -35/-10 boxes fused with a luciferase reporter gene (*luc*) (pJJ51), one carrying only the second -35/-10 promoter sequence fused with *luc* (pJJ52), and one carrying a promoterless *luc* gene (pJJ50). Both the luciferase activity and qRT-PCR results showed that *B. burgdorferi* strains harboring pJJ52 (only second -/35-10 promoter) or pJJ50 (no promoter) virtually had no detectable level of activity, whereas the strain carrying pJJ51 had over 100 fold higher luciferase activity and transcript levels than that of the promoterless control (**Fig. 1B**). These data confirmed the previous finding that the first -35/-10 promoter is the functional *glp* promoter, providing further genetic evidence for the presence of the long UTR in the *glp* operon.

**Fig. 1.**
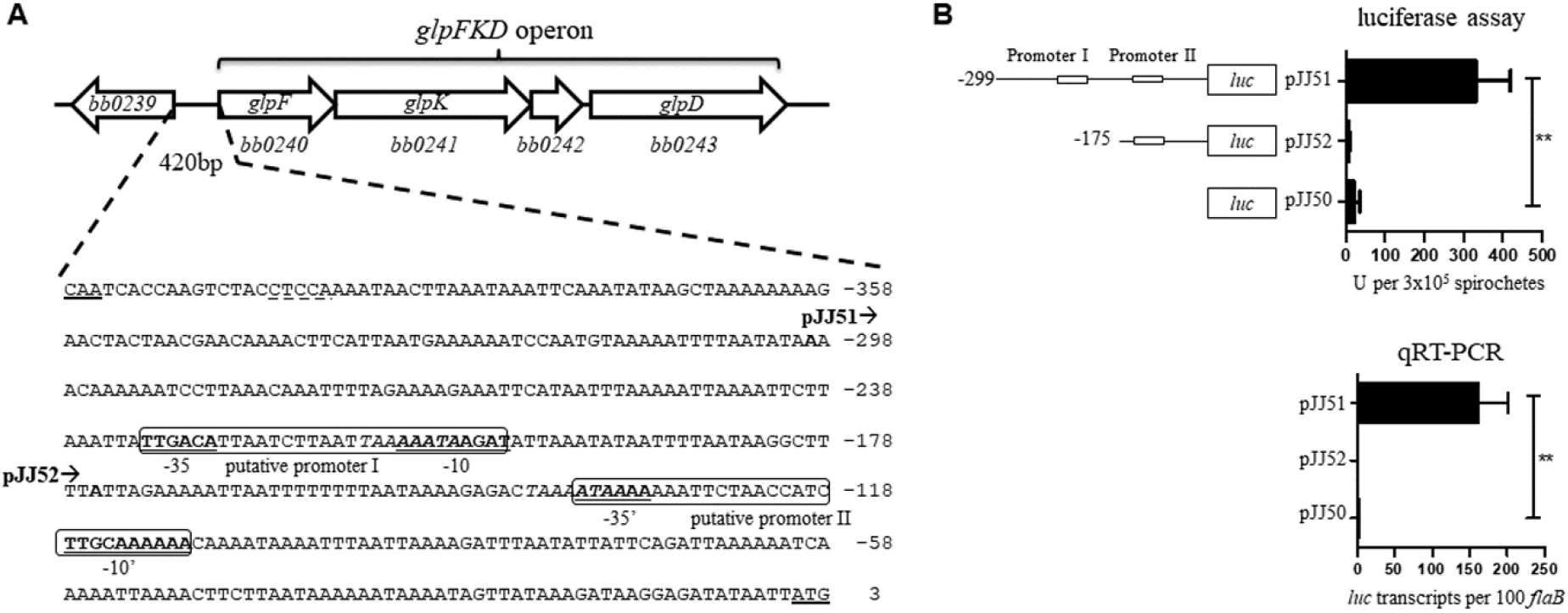
Prediction and genetic identification of the *glpFKD* promoter. (**A**) Schematic of the *glpFKD* operon (upper panel) and sequence of its upstream region (lower panel). The two predicted promoter regions of *glpFKD* operon (I and II) are boxed with -35/-10 motifs bolded and underlined. The translational start sites (TSS) are denoted by double underlines for the oppositely transcribed genes, *bb0240* (*glpF)*, and *bb0239*. The bold letters and arrows indicate the start positions and transcriptional direction of the promoter fragments for pJJ51 and pJJ52. The plasmids were transformed into *Borrelia* strain 5A4NP1. (**B**) Luciferase assay. Strains were grown in BSK-II medium and harvested (Mid log or stat, temp) for luciferase assay and qRT-PCR analysis. Results presented are means of 3 independent experiments performed in triplicate (± standard deviations). ** *p*<0.01 by student *t* test.

We next examined whether the UTR region is involved in regulating *glp* expression by constructing a series of truncated UTR fused with *luc* (**Fig. 2**, left). Deleting the entire UTR (pJJ56, denoted as minimal protomer P*glp*_min_) resulted in a 7-fold increase of luciferase activity relative to the full-length promoter (P*glp*, pJJ55) (**Fig. 2**, right), indicating that this UTR region plays a negative role in regulating *glp* expression. Deleting part of the UTR, the region I of the UTR (pJJ57), did not affect the promoter activity, suggesting that region II (−194 to -101) of UTR is involved in the negative regulation of *glp* expression. Replacing the RBS sequence of *glp* with that from the native luciferase gene (construct pJJ59) also did not affect luciferase activity. qRT-PCR analysis was consistent with the luciferase results (data not shown). Thus, negative regulation of *glp* by region II was at the transcriptional level.

**Fig. 2.**
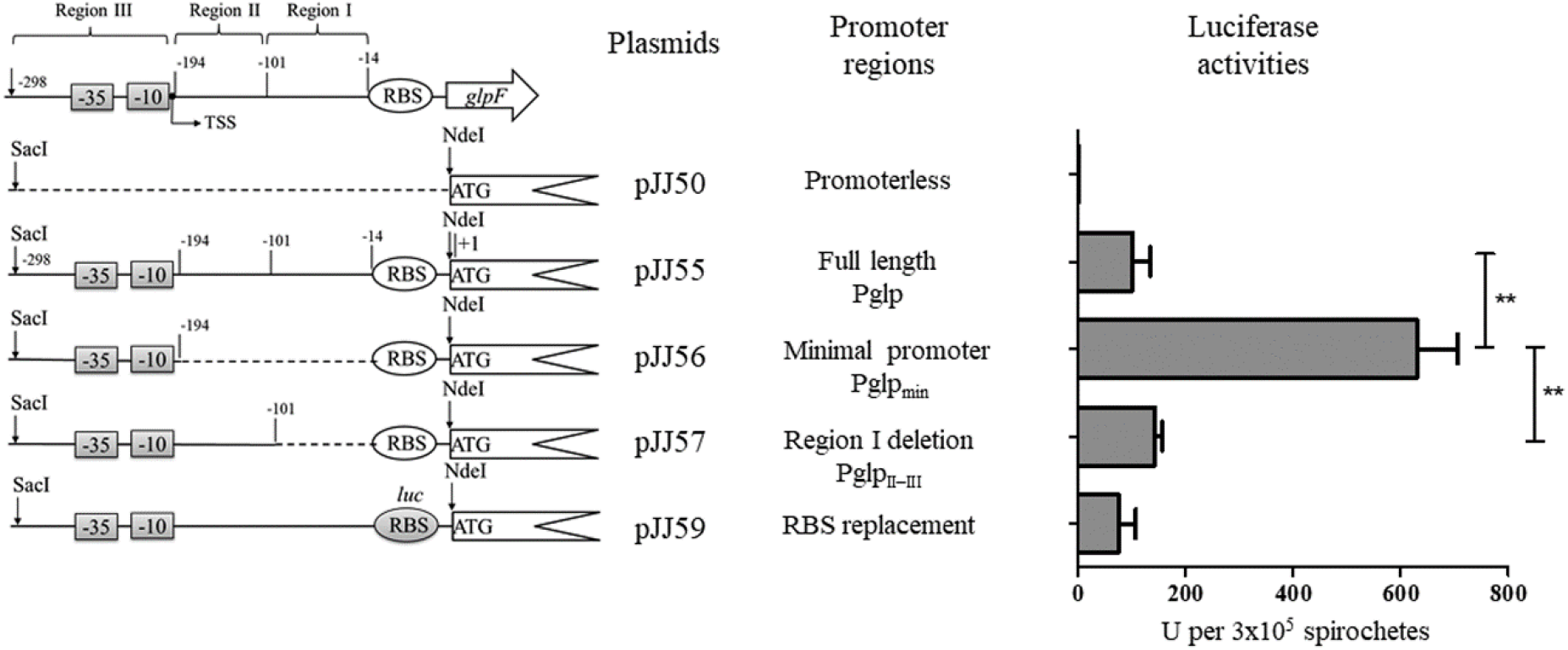
Identification of the *cis* element involved in *glpFKD* repression. Schematic showing various truncations of the *glpFKD* 5’ UTR fused with a luciferase reporter gene (*luc*). The -35 and -10 elements of *glp* operon are boxed with the putative transcriptional start site (TSS) indicated by an arrow (left panel). The construct names are labeled on the right with the description of their promoter composition. pJJ50, the plasmid containing a promoter less *luc* coding region; pJJ55, a full-length 5’ UTR of *glpF* fused with *luc*; pJJ56, a minimum promoter with complete deletion of 5’ UTR; pJJ57, deletion of the region I of 5’ UTR; pJJ59, identical to pJJ55 excepts that the *glpFKD* RBS sequence was replaced with the native RBS of *luc* gene. All constructs were transformed into wild-type *B. burgdorferi* strain 5A4NP1. The transformants were cultured in BSK-II medium and harvested at mid-log phase for luciferase assay (right panel) to quantify the promoter efficiency of each construct. Results presented are means of 3 independent experiments performed in triplicate (± standard deviations). ** *p*<0.01 by student *t* test.

### Identification of BadR as a repressor of *glp* expression

A common mechanism for a *cis* element to negatively regulate gene expression is to function as a repressor binding site. To test whether there is a repressor involved in regulation of *glp* expression, we overexpressed 13 predicted DNA-binding proteins present in the *B. burgdorferi* genome whose function are less known, including BB0253, BB0265, BB0345, BB0355, BB0361, BB0362, BB0364, BB0375, BB0434, BB0693, BB0805, BB0831, and BBD22. Overexpression of one of these proteins, BB0693 (BadR), significantly repressed *glp* expression (**Fig 3A**). Given that BadR (Borrelia Host Adaptation Regulator) was previously shown to repress *rpoS* and *bosR* expression (26, 27), we then examined the effect of *badR* over expression and deletion on *glp* expression in *B. burgdorferi*. Consistent with our previous findings, *badR* over expression resulted in significant reduction in glp expression (**Fig. 3B**). Though,the *rrp1* mutant had dramatically reduced *glp* expression as we previously shown (20), the *badR* mutant showed a 7-fold increase of *glpF* expression compared to wild-type *B. burgdorferi* **(Fig. 3C)**. Thus, overexpression and deletion of *badR* results strongly suggest that BadR is a repressor of *glp*.

**Fig. 3.**
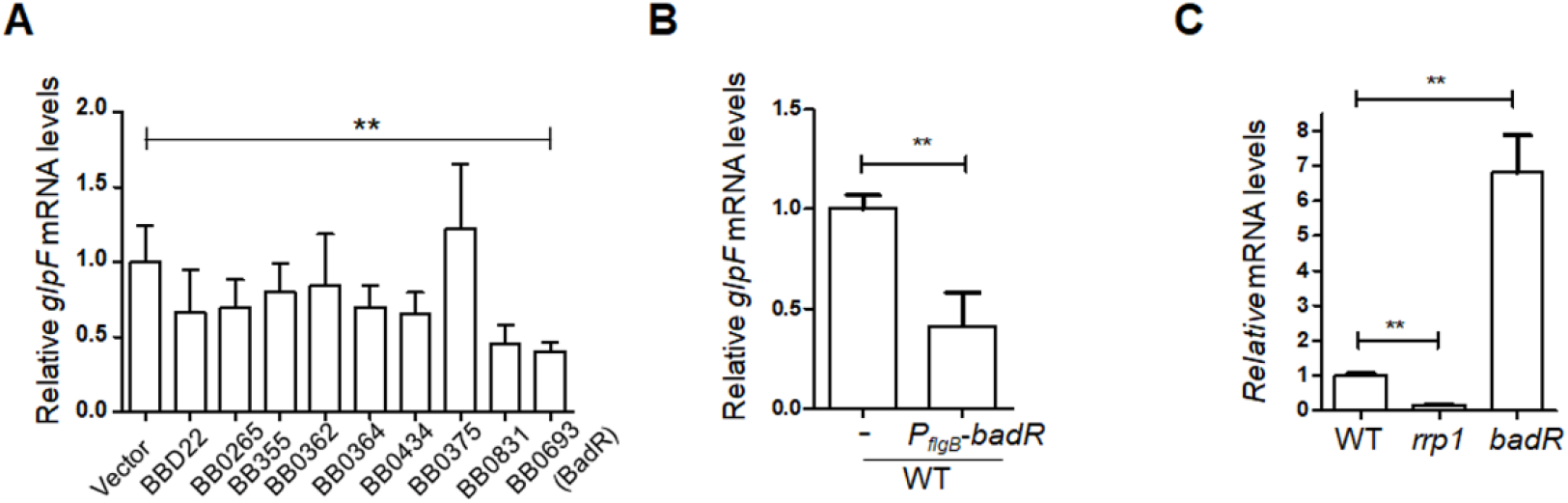
BadR is a repressor of the *glpFKD* operon. **(A)** Effect of inducible expression of various *Bb* transcriptional regulatory proteins on *glpF*. Relative transcript levels of the glycerol operon *glpFKD* (represented by *glpF*) were determined in the wild-type (vector along) and Borrelia harbouring IPTG-driven transctiptional regulators (as mentioned in the legend), (**B**), Relative transcript levels of the glycerol operon *glpFKD* across wt and *badR* over expressing strains. (C) Relative transcript levels **of *glpF*** in *rrp1* or *badR* mutant. For all the expreriments, RNA was isolated from mid-logarithmic phase cultures grown at 37°C in standard BSK-II medium. The levels of *glpF* expression in the wild-type *Borrelia* were set as a value of 1, and relative levels of *glpF* in each sample were reported. Results presented are means of 3 independent experiments performed in triplicate (± standard deviations). ** *p*<0.01 by student *t* test.

### BadR binds to the *glp* promoter

To gather biochemical evidence that BadR is the repressor of *glp*, electrophoretic mobility shift assays (EMSAs) were performed using various amounts of recombinant BadR with ^32^P-labeled DNA fragments containing various lengths of *glp* promoter (**Fig. 4A**). The assay was conducted in the presence of 1 μg of dI: dC as a nonspecific competitor. The result showed that BadR binds to the DNA fragment containing the full-length promoter with the complete UTR (Pglp I-II-III), or the promoter with part of UTR containing region II (Pglp II-III), but not region II alone (PglpII), or the upstream fragment region III alone (Pglp III) (**Fig. 4B**, top panel), suggesting that BadR binds to the *glp* promoter, in particular, region II-III surrounding the TSS site. The binding of BadR to the labeled DNA probes was inhibited in the presence of an excess of unlabeled DNA, and no binding was observed when using a DNA fragment containing an unrelated *rrp2* sequence (data not shown), suggesting that the BadR binding to the *glp* promoter is specific.

**Fig. 4.**
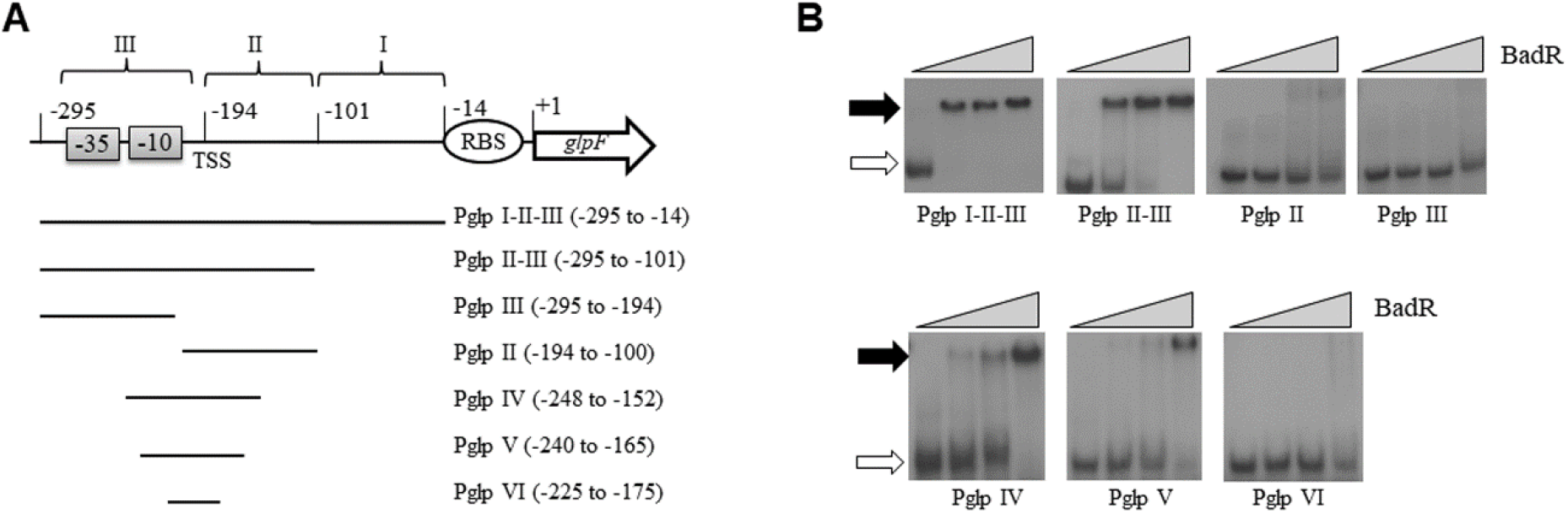
BadR binds directly to the *cis* regulatory region for the modulation of *glp* expression. The electrophoretic mobility shift assays (EMSAs) were conducted using purified BadR and various DNA fragments upstream of *glpF*. (**A**) The lengths of different DNA probes and their relative positions to the *glp* operon are indicated. The ATG of *glpF* as set as +1. Various amounts of purified BadR (0, 0.5, 1.0, 2.0 pmol) were incubated with 0.2 pmol of ^32^P labeled DNA in 10 μl reactions for 30 min at 23°C before subjected to non-denatured polyacrylamide gels (7.5%) analysis. Gels were dried, and radioactive signals were visualized by exposure to X-ray film. (**B**) Probe used in the assay is specified below the image. The free probes are indicated by open arrows and the retarded DNA-protein complex is represented by solid arrows.

To further identify the region required for BadR binding, the region II-III was further truncated. BadR was capable of binding to a 100 bp DNA fragment with 50 bp upstream and downstream of TSS (Pglp IV), or an 80 bp DNA fragment with 40 bp upstream and downstream of TSS (Pglp V) (**Fig. 4B**, bottom panel), but not 50 bp DNA fragment with 25 bp on both side of TSS (Pglp VI). These data suggest that the 80 bp region surrounding TSS (−240 to -165) is the core region for BadR binding.

### The binding of BadR to the *glp* promoter is not affected by sugars and phosphorylated sugars

BosR belongs to the ROK (repressor, opening reading frame, kinase) protein family, which functions as carbohydrate-responsive transcriptional repressors that generally interact with various carbon sources (28, 29). Previously it was shown that BadR binds to the *rpoS* promoter and several phosphorylated sugars, such as glucose-6P, inhibit the binding (26). Given that BadR represses *glp* and glycerol dramatically induces *glp* expression (19), it is conceivable that glycerol, glycerol-3-phosphate, or phosphorylated sugars may affect BadR binding to *glp*. To test this possibility, EMSAs were carried out in the presence of various sugars or phosphorylated sugars. The assays were conducted in the presence of 0.7 pmol BadR with ^32^P-labeled 200 bp DNA fragment (PglpII-III). The results showed that the binding affinity of BadR to the *glp* operon was not affected by the presence of glycerol or glucose, nor by the presence of phosphorylated sugars, including glycerol-3-phosphate (Glycerol-3-P), glucose-6-phosphate (glucose-6-P), ribose-5-phosphate (Ribose-5-P) and N-acetylglucosamine-6-phosphate (GlcNAc-6-P) (**Fig. 5)**.

**Fig. 5.**
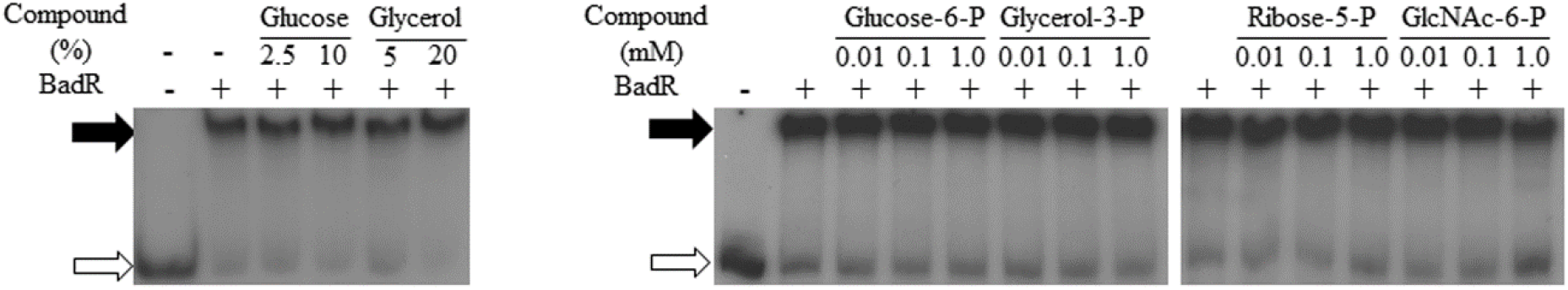
Binding of BadR to *glpFKD* operon is not affected by sugars or phosphorylated sugars. The effects of chemical and protein factors on the binding affinity of BadR with *glpFKD* operon were investigated by EMSAs as the method described in Fig. 4. The DNA fragment PglpII-III (as illustrated in Fig. 4) was labeled with ^32^P as the probe (0.2 pmol). The total reaction volume was 10 μl containing 0.7 pmol purified BadR. Various amounts of sugars and phosphorylated sugars were added to the reaction as indicated. The free probes are indicated by open arrows and the retarded DNA fragments by solid arrows.

## DISCUSSION

Previous studies established the importance of the *glp* operon for *B. burgdorferi*, including its role in persistence within ticks and transmission to mice (4, 13-15, 20). The spirochete shifts its carbohydrate utilization as it traverses the enzootic cycle, favoring glucose in the vertebrate host and glycerol (along with chitobiose) in the tick vector. However, little is known about how *B. burgdorferi* modulates its carbohydrate metabolism. In this study, for the first time, we identified a direct regulator for the modulation of *glp* operon expression in *B. burgdorferi*. We provided both genetic and biochemical evidence to reveal that BadR represses transcription of the *glpFKD* operon in *B. burgdorferi* by binding to an 80 bp fragment flanking TSS. BadR is then proposed to govern the catabolic switch, allowing appropriate adaptation to the challenges of a dynamic and hostile environment in tick midgut for *B. burgdorferi*.

BadR (*<u>B</u>orrelia* host <u>ad</u>aptation <u>r</u>egulator) was discovered by Seshu’s group in 2013 as a repressor that modulates host adaptation and virulence in *B. burgdorferi* (26). It belongs to the ROK family of proteins with DNA-binding ability and is a presumed regulator that responds to sugar(s) or its(their) metabolic intermediate(s) (28). Transcriptomic studies demonstrated that BadR is a global regulator in *B. burgdorferi* (26) (30). RpoS, BosR, and *Borrelia* host adaptation protein (BadP) were the direct targets of BadR identified so far (26, 27, 31). In this study, we further demonstrated that BadR can also directly modulate glycerol utilization by *B. burgdorferi* in response to glycerol availability. The function of BadR in the glycerol metabolism added additional evidence for the diversity of ROK family, besides NagC and Mlc from *E. coli*, XylR from *Bacillus subtilis* and CsnR from *Streptomyces lividans* which are involved in the utilization of N-acetylglucosamine (GlcNAc) (32), xylose (33) and chitosanase (34), respectively.

The signal that triggers the release of BadR from its targets is still unclear. There are controversial reports for BadR effectors or inducers in *B. burgdorferi*. Miller et al. reported that various phosphorylated sugars affected the binding affinity of BadR to the *rpoS* promoter (26), while Ouyang et al. found an opposite phenomenon (27). Our data suggested that the binding of BadR to the *glp* operon was not affected by glucose, glycerol, ribose, GlcNac, or phosphorylated sugars (**Fig. 5**). The regulation of ROKs generally involves interaction with their inducing signals, some unknown *Borrelia*-specific metabolic intermediate(s) may act to modulate the DNA-binding affinity of BadR in *Borrelia*. Meanwhile, the effector specificity-determining residues for ROKs are not well conserved in BadR (27, 29), implying different induction mechanisms for the regulatory complex of BadR-glycerol-*glp* operon.

The *glp* operon has an unexpectedly long 5’ UTR (∼200 nts) comparing the median 5’UTR length in *B. burgdorferi* (∼36 nts) (13, 35) (**Fig. 1A**). The locations of the *glp* operon and its Pribnow box identified in the current study were consistent with a global mapping study of the *Borrelia* transcriptome by RNA-seq (25) and 5’ rapid amplification of cDNA ends of *glpFKD* operon (35). We further characterized the upstream region of the *glp* operon and identified an 80 bp core fragment crossing the TSS as the *cis*-element for *glp* regulation by both promoter-luciferase reporter assay and BadR: DNA gel shift assay. Promoter activity assay indicated that the deletion of the 5’ UTR increased the *glp* promoter activity by 7-fold (JJ56 vs. JJ55, Fig. 2) in the wild-type *Borrelia*. Consistently, a 7-fold upregulation of *glpF* transcription was observed in the *badR* mutant (**Fig. 3C**). These demonstrated that BadR repressed *glp* expression by binding with the *cis* element in *Borrelia*.

The structure and regulation of the *glp* operon are remarkably more complex than previously appreciated. Multiple canonical intracellular signaling pathways, including RpoS, c-di-GMP, and (p)ppGpp, converge on the tight control of *glp* expression (4, 11, 14, 16, 20-22). Besides these global regulators, SpoVG may be another direct regulator since it can bind multiple sites with both DNA and mRNA of the *glp* operon (36). Moreover, *glpD* (*bb0243*), the last gene in this operon, is differentially regulated with *glpF* and *glpK* in multiple transcriptome-based analyses (15, 16, 25). A *cis*-regulatory RNA element was identified to be associated with the 5’ UTR region of *glp* operon (25). These studies suggested the presence of post-transcriptional regulation for glycerol utilization in *Borrelia*. The interplay of multiple regulatory machines, therefore, controls the carbohydrate metabolism switch *in vivo* for the acclimation of *Borrelia* to its complicated life cycle. It will be interesting to investigate further how these different regulatory factors coordinate to reciprocally modulate glycerol gene expression throughout the tick-mouse enzootic cycle and tether regulatory networks to global cellular functions.

## MATERIALS AND METHODS

### Bacterial strains and growth conditions

The strains and plasmids used in this study are listed in **Table 1**. Low-passage, virulent *B. burgdorferi* strain 5A4NP1 is the wild-type strain used in this study. *Borrelia* strains were generally cultivated in BSK-II medium (37) with 6% rabbit serum at 37°C with 5% CO_2_. Relevant antibiotics were added to the cultures with the following concentrations when necessary, 300 μg ml^-1^ kanamycin, 100 μg ml^-1^ streptomycin, and 100 μg ml^-1^ gentamicin. The shuttle vector was constructed and maintained in *Escherichia coli* DH5α. The transformation of *Borrelia* and screening was conducted according to a previous report (38).

**Table 1.** Strains and plasmids used in this study.

| Strains and plasmids | Phenotypes | Source |
| --- | --- | --- |
| *Borrelia* |  |  |
| 5A4NP1 | Wild-type parental strain, low-passage and virulent strain derived from B31 | (9) |
| *rrp1* | 5A4NP1 with *rrp1* disrupted with an *aadA* marker conferring streptomycin resistance | (9) |
| *badR* | 5A4NP1 with *badR* disrupted with an *aaC1* marker conferring gentamicin resistance | This study |
| Plasmids |  |  |
| pET100/D-TOPO | Expression vector with an N-terminal His<sub>6</sub> tag | Invitrogen |
| pMAL-c2X | Expression vector fused with N-terminal MBP | NEB |
| pSC-A-amp/kan | TA cloning vector | Agilent |
| pGEM-T | TA cloning vector | Promega |
| pJD48 | pBSV2-based shuttle vector with promoterless *luc*<sub>Bb</sub><sup>+</sup> (*Borrelia* codon-optimized luciferase gene), Kan<sup>r</sup> | (40) |
| pJD50 | pJD48 with Erm substituted Kan resistance marker, Erm<sup>r</sup> |  |
| pBT50 | pJD50 with promoterless *luc*<sub>Bb</sub><sup>+</sup>, Erm<sup>r</sup> | This study |
| pJJ50 | pBT50 with the replacement of Erm with Gen resistance marker, Gen<sup>r</sup> | This study |
| pJJ51 | pJJ50 with *glpFKD* putative promoter I-driven *luc*<sub>Bb</sub><sup>+</sup> with a *luc* native RBS. | This study |
| pJJ52 | pJJ50 with *glpFKD* putative promoter II-driven *luc*<sub>Bb</sub><sup>+</sup> with a *luc* native RBS. | This study |
| pJJ55 | pJJ50 with *glpFKD* promoter-driven *luc*<sub>Bb</sub><sup>+</sup> | This study |
| pJJ56 | pJJ50 with the deletion of 5' UTR of *glpFKD* promoter | This study |
| pJJ57 | pJJ50 with the deletion of the downstream of 5' UTR of *glpFKD* promoter | This study |
| pJJ59 | pJJ55 with the replacement of *glpFKD* RBS with native luciferase RBS | This study |
| pMH105 | Source of gentamicin resistance marker (*PflaB-aaC1*) | (20, 44) |
| pJSB275 | Shuttle vector with IPTG-inducible luciferase gene | (41, 45) |
| pJJ275R | Originated from pJSB275 with the replacement of luciferase gene fragment (NdeI-HindIII) with multiple cloning sites encoding N-terminal FLAG and C-terminal HA tags | This study |
| pJD55 | pBSV2-derived shuttle vector with *flaB* promoter driven *rpoN* | (46, 47) |
| pJJ43 | pJD55 with *PflgB-bb0693* replacing *PflaB-rpoN* | This study |
| pJJ49 | For the creation of bb0693 knockouts originated from pSC-A-amp/kan | This study |
| pJJ53 | *bb0693* cloned into pMAL-c2X to express soluble Bb0693 | This study |

### RNA extraction and qRT-PCR

RNA samples were extracted from mid-log phase *B. burgdorferi* cultures (around 10^8^ cells) using the RNeasy Mini kit (Qiagen). DNA contamination in the RNA samples was removed by DNase I (New England Biolabs, NEB) digestions, and confirmed by PCR amplifying the *flaB* gene of *B. burgdorferi*. Three independent culture samples were prepared for each strain. The cDNA was prepared from 1 μg RNA using Superscript III Reverse Transcriptase with random primers (Invitrogen). qRT-PCR was performed using SYBR Green PCR Master Mix (Applied Biosystems) on an ABI 7000 sequence detection system. The *flaB* gene of *B. burgdorferi* was used as a reference. The quantities of the targeted genes and *flaB* in cDNA samples were calculated according to our previous report (9). The relative transcript level was determined by the threshold cycle (2^*-ΔΔCt*^) method (39). An unpaired *t-*test was conducted to determine the statistical significance of different values in the qRT-PCR results.

### *B. burgdorferi glpFKD* -luciferase reporter constructs and luciferase assay

The *Borrelia* codon-optimized luciferase gene cassette (*luc*_*Bb+*_) from pJD48 was first digested by AvaI and BglII (40) and ligated into plasmid pJD50 (an erythromycin-resistant derivative of the borrelial shuttle vector pBSV2) to generate pBT50. To replace the erythromycin resistance marker of pBT50, the gentamicin resistance marker (P_*flaB*_-*aaC1*) was amplified from pMH105 (20) by PCR and then TA-cloned into pSC-A-amp/kan (Agilent Technologies). Following confirmation of the TA clones, the P_*flaB*_-*aaC1* fragment was digested by SacI and BspHI and cloned into pBT50 linearized by digestion with the same enzymes. The resulting plasmid, designated as pJJ50, was used as a promoterless control in the following promoter-luciferase reporter system. To verify the activities of the two putative promoters (I and II as indicated in **Fig. 1A**), the 299 bp and 175 bp fragments upstream of the translation start site of *glpF* were amplified by PCR with the genomic DNA of the wild-type *Borrelia* as the template, using pair of primers that were engineered to contain a SacI (forward primer) or an NdeI site (reverse primer) at their 5’ ends. PCR products were double-digested with SacI-NdeI and cloned into SacI-NdeI-digested pJJ50 to generate plasmids pJJ51 and pJJ52, respectively. The RBS from the native luciferase gene was used in the above two plasmids. A similar strategy was used to generate *glp* -luciferase reporter plasmids, including pJJ55, pJJ56, pJJ57, and pJJ59 (**Fig. 2**). The luciferase gene in all other constructs was driven by the native *glp* promoter with truncated *glp* 5’ UTR. For the construction of pJJ55, the DNA fragment upstream of *glpF* containing the full length of 5’UTR and promoter region of *glp* operon were amplified by PCR and then cloned into the shuttle plasmid pJJ50. The plasmid, pJJ59, was constructed by substituting the *glp* operon’s Shine-Dalgarno Sequence (RBS) in pJJ55 with that from the luciferase gene. All the above shuttle plasmids were sequenced before they were transformed into *Borrelia* strains. All *Borrelia* strains carrying the testing constructs were inoculated into fresh BSK-II medium for growth to mid-log phase (3×10^6^ to 3×10^7^ cells ml^-1^), and then two 5 ml aliquots were collected by centrifugation. One aliquot was used for RNA extraction and qRT-PCR analysis, and the other one was lysed for luciferase assay (40) using a commercial luciferase system (Promega). The luciferase activity in the test tubes was measured using a Turner BioSystems TD-20/20 Luminometer and reported as relative luciferase units (RLU) with average background luminescence subtracted from readings.

### Screening of the potential *glpFKD* expression repressors

The shuttle plasmid pJJ275R was constructed based on the modification of pJSB275 (41) to facilitate IPTG-inducible gene expression in *Borrelia*. For this purpose, pJSB275 was digested with NdeI and HindIII to excise the luciferase gene and insert a synthetic DNA fragment with multiple cloning sites encoding N-terminal FLAG and C-terminal HA tags. Several putative transcription factors or proteins with DNA-binding domains were selected for *glpFKD* regulator screening. These candidates include BBD22, BB0253, BB0265, BB0345, BB0355, BB0361, BB0362, BB0364, BB0375, BB0434, BB0805, BB0831 and BB0693. The DNA fragments carrying the candidate genes were cloned into the NdeI and XhoI sites of pJJ275R, in which they were fused with an N-terminal FLAG tag controlled by the promoter of pQE30. The resulting plasmids were transformed into *Borrelia*. Expression of the gene of interest is induced with the addition of IPTG (1 mM), which can be confirmed by immunoblot analysis using anti-FLAG antibodies. To quantify the *glpFKD* expression levels, 10 ml *Borrelia* cultures were grown to the mid-log phase in the presence of 1 mM IPTG with pJJ275R as a control. The cells were then harvested for RNA extraction and qRT-PCR analysis.

### Construction of *bb0693* (*badR*) overexpression *Borrelia* strain

A pBSV2-derived shuttle vector, pJD55 (40), was used to constitutively overexpress *bb0693* in *B. burgdorferi*. The gene fragments containing *bb0693* and the *flgB* promoter region were amplified and fused by overlap PCR. The restriction sites, BglII and SmaI, were incorporated into the designated primers for the insertion of the PCR fragment into pJD55 to generate pJJ43. Thus, the *bb0693* gene was driven by a strong promoter in the shuttle vector and encoded a C-terminal HA-tagged protein. The resulting plasmid was verified by sequencing before being transformed into *Borrelia*. The gentamycin resistance transformants were verified by immunoblot analysis using the anti-HA monoclonal antibody to confirm the constitutive overexpression of *bb0693*.

### Inactivation of *badR* in *B. burgdorferi*

The *badR* mutant has been published by Seshu’s group and Ouyang’s group previously. A suicide vector, pJJ49, was constructed for the disruption of the *badR* gene through an allelic exchange. Briefly, around 1.4 kb regions of the *Borrelia* chromosome both upstream and downstream of the *badR* gene were PCR amplified from the total genomic DNA of the wild-type strain 5A4NP1. The resulting DNA fragments were then cloned into a gentamicin-resistant marker (*aaC1*) driven by a constituently expressed *Borrelia flaB* promoter via PstI/BamHI and MluI/SacI. The resulting plasmid, pJJ49, was confirmed by both restriction enzyme digestion and sequencing, followed by *Borrelia* transformation. The transformants were screened in the presence of gentamicin and kanamycin in the BSK-II medium. The correct marker insertion and *badR* inactivation were verified by PCR, and positive clones were further confirmed for the loss of BadR by immunoblot analysis and qRT-PCR. Plasmid profiles of the confirmed *badR* clones were determined by multiple PCR analyses with nineteen pairs of primers specific for each of the endogenous plasmids as reported (42). One of the *badR* mutant clones that had plasmid profiles identical to the parental strain was chosen for further study.

### Overexpression and purification of BadR

The full-length *badR* gene was amplified from the genome DNA of the wild-type *Borrelia* using High-Fidelity Taq Polymerase Fusion (NEB). The fragment was subsequently ligated into the pMAL-c2X (NEB) expression vector via BamHI/SalI, which codes for an N-terminal maltose-binding protein fused with BadR. The resulting plasmid, pJJ53, was transformed into Rosetta™ (Novagen) cells after confirmation by restriction enzyme digestion analysis and sequencing. The BadR protein was solubly expressed when cells were induced overnight at 16°C with 0.4 mM IPTG after being grown to an optical density of 0.6 at 600 nm in Lysogeny broth (LB) medium. The cells were collected by centrifugation, and protein purification was carried out according to the manual of Amylose Resin (NEB). The protein, as well as the purification procedure, was monitored by sodium dodecyl sulfate-polyacrylamide gel electrophoresis (SDS-PAGE) to confirm that the proteins were purified to homogeneity. Proteins were stored at -80°C, and concentrations were determined using the Bio-Rad protein assay with BSA as a standard (Bio-Rad).

### EMSAs

EMSAs were performed to determine protein: DNA interactions as described previously (43). The *glpFKD* promoter fragments were labeled with ^32^P using DNA Polymerase I Large Fragment (Klenow fragment) (NEB) to fill in the 5’ overhang with [α-^32^P]dATP. Labeled DNA fragments (0.2 pmol) and various amounts of purified BadR were mixed in 10 μl binding buffer in the presence of 100 μg ml^-1^ poly dI: dC and incubated for 30 min at 23°C. The reactions were then analyzed on 7.5 % non-denatured polyacrylamide gels and the protein: DNA complex was detected as a retarded band on the gel. Some reactions included unlabeled promoter competitors or chemicals for competition studies. Gels were dried and exposed in a cassette using an X-ray film for autoradiography.

### SDS-PAGE and immunoblot analysis

*B. burgdorferi* strain was inoculated into BSK-II medium and grown to a density of approximately 3×10^7^ cells ml^-1^ before it was harvested by centrifugation at 7,000 g and washed twice with phosphate buffered saline (PBS, 50 mM, pH7.4). The pellets were collected for SDS-PAGE and immune blotting. Immunoblots were developed using the SuperSignal West Pico chemiluminescent substrate according to the manufacturer’s instructions (Pierce). FLAG-tagged or HA-tagged protein was detected using a commercially available mouse monoclonal antibody (Promega).

## ACKNOWLEDGMENTS

Funding for this work was partially provided by NIH grants AI083640 (to X.F.Y.) and the National Science Foundation of China 81772229 and support of Key Discipline of Zhejiang Province in Medical Technology (First Class, Category A) (to Y.L). This investigation was partially conducted in a facility with support from research facilities improvement program grant number C06 RR015481-01 from the National Center for Research Resources, NIH.

## REFERENCES

1. Radolf JD, Caimano MJ, Stevenson B, Hu LT. 2012. Of ticks, mice and men: understanding the dual-host lifestyle of Lyme disease spirochaetes. Nat Rev Micro 10:87–99.

2. Steere AC, Grodzicki RL, Kornblatt AN, Craft JE, Barbour AG, Burgdorfer W, Schmid GP, Johnson E, Malawista SE. 1983. The spirochetal etiology of Lyme disease. The New England Journal of Medicine 308:733–740.

3. Fraser CM, Casjens S, Huang WM, Sutton GG, Clayton R, Lathigra R, White O, Ketchum KA, Dodson R, Hickey EK, Gwinn M, Dougherty B, Tomb JF, Fleischmann RD, Richardson D, Peterson J, Kerlavage AR, Quackenbush J, Salzberg S, Hanson M, van Vugt R, Palmer N, Adams MD, Gocayne J, Weidman J, Utterback T, Watthey L, McDonald L, Artiach P, Bowman C, Garland S, Fujii C, Cotton MD, Horst K, Roberts K, Hatch B, Smith HO, Venter JC. 1997. Genomic sequence of a Lyme disease spirochaete, Borrelia burgdorferi. Nature 390:580–586.

4. Corona A, Schwartz I. 2015. Borrelia burgdorferi: Carbon metabolism and the Tick-Mammal enzootic cycle. Microbiol Spectr 3.

5. Casjens S, Palmer N, van Vugt R, Huang WM, Stevenson B, Rosa P, Lathigra R, Sutton g, Peterson J, Dodson RJ, Haft D, Hickey E, Gwinn M, White O, Fraser CM. 2000. A bacterial genome in flux: the twelve linear and nine circular extrachromosomal DNAs in an infectious isolate of the Lyme disease spirochete Borrelia burgdorferi. Molecular Microbiology 35:490–516.

6. Gherardini F, Boylan J, Lawence K, Skare J. 2010. Metabolism and Physiology of Borrelia, p 103-138. In Samules DS, Radolf JD (ed), Borrelia: Molecular Biology, Host Interaction and Pathogenesis. Caister Academic Press, Norfolk, UK.

7. de Silva AM, Fikrig E. 1995. Growth and migration of Borrelia burgdorferi in Ixodes ticks during blood feeding. American Journal of Tropical Medicine and Hygiene 53:397–404.

8. Dunham-Ems SM, Caimano MJ, Pal U, Wolgemuth CW, Eggers CH, Balic A, Radolf JD. 2009. Live imaging reveals a biphasic mode of dissemination of Borrelia burgdorferi within ticks. J Clin Invest 119:3652–65.

9. He M, Ouyang Z, Troxell B, Xu H, Moh A, Piesman J, Norgard MV, Gomelsky M, Yang XF. 2011. Cyclic di-GMP is essential for the survival of the Lyme disease spirochete in ticks. PLoS Pathog 7:e1002133.

10. Sze CW, Smith A, Choi YH, Yang X, Pal U, Yu A, Li C. 2013. Study of the response regulator Rrp1 reveals its regulatory role in chitobiose utilization and virulence of Borrelia burgdorferi. Infect Immun 81:1775–1787.

11. Caimano MJ, Dunham-Ems S, Allard AM, Cassera MB, Kenedy M, Radolf JD. 2015. Cyclic di-GMP modulates gene expression in Lyme disease spirochetes at the tick-mammal interface to promote spirochete survival during the blood meal and tick-to-mammal transmission. Infect Immun 83:3043–60.

12. Vandyk JK, Bartholomew DM, Rowley WA, Platt KB. 1996. Survival of Ixodes scapularis (Acari: Ixodidae) exposed to cold. J Med Entomol 33:6–10.

13. Pappas CJ, Iyer R, Petzke MM, Caimano MJ, Radolf JD, Schwartz I. 2011. Borrelia burgdorferi requires glycerol for maximum fitness during the tick phase of the enzootic cycle. PLoS Pathog 7:e1002102.

14. Caimano MJ, Drecktrah D, Kung F, Samuels DS. 2016. Interaction of the Lyme disease spirochete with its tick vector. Cellular Microbiology 18:919–927.

15. Iyer R, Caimano MJ, Luthra A, Axline D, Corona A, Iacobas DA, Radolf JD, Schwartz I. 2015. Stage - specific global alterations in the transcriptomes of Lyme disease spirochetes during tick feeding and following mammalian host adaptation. Molecular microbiology 95:509–538.

16. Drecktrah D, Lybecker M, Popitsch N, Rescheneder P, Hall LS, Samuels DS. 2015. The Borrelia burgdorferi RelA/SpoT homolog and stringent response regulate survival in the tick vector and global gene expression during starvation. PLoS Pathog 11:e1005160.

17. Caimano MJ, Kenedy MR, Kairu T, Desrosiers DC, Harman M, Dunham-Ems S, Akins DR, Pal U, Radolf JD. 2011. The hybrid histidine kinase Hk1 is part of a two-component system that is essential for survival of Borrelia burgdorferi in feeding Ixodes scapularis ticks. Infect Immun 79:3117–3130.

18. Kostick JL, Szkotnicki LT, Rogers EA, Bocci P, Raffaelli N, Marconi RT. 2011. The diguanylate cyclase, Rrp1, regulates critical steps in the enzootic cycle of the Lyme disease spirochetes. Mol Microbiol 81:219–231.

19. Zhang J-J, Chen T, Yang Y, Du J, Li H, Troxell B, He M, Carrasco SE, Gomelsky M, Yang XF. 2018. Positive and Negative Regulation of Glycerol Utilization by the c-di-GMP Binding Protein PlzA in Borrelia burgdorferi. Journal of Bacteriology 200:e00243–18.

20. He M, Zhang J-J, Ye M, Lou Y, Yang XF. 2014. The cyclic di-GMP receptor PlzA controls virulence gene expression through RpoS in Borrelia burgdorferi. Infect Immun 82:445–52.

21. Caimano M, Iyer R, Eggers C, Gonzalez C, Morton E, Gilbert M, Schwartz I, Radolf J. 2007. Analysis of the RpoS regulon in Borrelia burgdorferi in response to mammalian host signals provides insight into RpoS function during the enzootic cycle. Mol Microbiol 65:1193 – 1217.

22. Bugrysheva JV, Pappas CJ, Terekhova DA, Iyer R, Godfrey HP, Schwartz I, Cabello FC. 2015. Characterization of the Rel_Bbu_ regulon in Borrelia burgdorferi reveals modulation of glycerol metabolism by (p) ppGpp. PloS one 10:e0118063.

23. Rogers EA, Terekhova D, Zhang H, Hovis KM, Schwartz I, Marconi RT. 2009. Rrp1, a cyclic-di-GMP-producing response regulator, is an important regulator of Borrelia burgdorferi core cellular functions. Mol Microbiol 71:1551–73.

24. Grove AP, Liveris D, Iyer R, Petzke M, Rudman J, Caimano MJ, Radolf JD, Schwartz I. 2017. Two distinct mechanisms govern RpoS-mediated repression of tick-phase genes during mammalian host adaptation by Borrelia burgdorferi, the Lyme Disease spirochete. mBio 8:e01204–17.

25. Adams PP, Flores Avile C, Popitsch N, Bilusic I, Schroeder R, Lybecker M, Jewett MW. 2016. In vivo expression technology and 5′ end mapping of the Borrelia burgdorferi transcriptome identify novel RNAs expressed during mammalian infection. Nucleic Acids Research doi:10.1093/nar/gkw1180.

26. Miller CL, Karna SLR, Seshu J. 2013. Borrelia host adaptation regulator (BadR) regulates rpoS to modulate host adaptation and virulence factors in Borrelia burgdorferi. Mol Microbiol 88:105–24.

27. Ouyang Z, Zhou J. 2015. BadR (BB0693) controls growth phase-dependent induction of rpoS and bosR in Borrelia burgdorferi via recognizing TAAAATAT motifs. Mol Microbiol 98:1147–1167.

28. Titgemeyer F, Reizer J, Reizer A, Saier MH, Jr. 1994. Evolutionary relationships between sugar kinases and transcriptional repressors in bacteria. Microbiology 140 (Pt 9):2349–54.

29. Kazanov MD, Li X, Gelfand MS, Osterman AL, Rodionov DA. 2012. Functional diversification of ROK-family transcriptional regulators of sugar catabolism in the Thermotogae phylum. Nucleic Acids Research 41:790–803.

30. Arnold WK, Savage CR, Lethbridge KG, Smith TC, 2nd, Brissette CA, Seshu J, Stevenson B. 2018. Transcriptomic insights on the virulence-controlling CsrA, BadR, RpoN, and RpoS regulatory networks in the Lyme disease spirochete. PLOS ONE 13:e0203286.

31. Smith TC, Helm SM, Chen Y, Lin Y-H, Rajasekhar Karna SL, Seshu J. 2018. Borrelia host adaptation protein (BadP) Is required for the colonization of a mammalian host by the agent of Lyme disease. Infection and Immunity 86:e00057–18.

32. Plumbridge J. 2001. Regulation of PTS gene expression by the homologous transcriptional regulators, Mlc and NagC, in Escherichia coli (or how two similar repressors can behave differently). J Mol Microbiol Biotechnol 3:371–80.

33. Dahl MK, Degenkolb J, Hillen W. 1994. Transcription of the xyl operon is controlled in Bacillus subtilis by tandem overlapping operators spaced by four base-pairs. J Mol Biol 243:413–24.

34. Dubeau MP, Poulin-Laprade D, Ghinet MG, Brzezinski R. 2011. Properties of CsnR, the transcriptional repressor of the chitosanase gene, csnA, of Streptomyces lividans. J Bacteriol 193:2441–50.

35. Grove AP, Liveris D, Iyer R, Petzke M, Rudman J, Caimano MJ, Radolf JD, Schwartz I. 2017. Two Distinct Mechanisms Govern RpoS-Mediated Repression of Tick-Phase Genes during Mammalian Host Adaptation by Borrelia burgdorferi, the Lyme Disease Spirochete. MBio 8.

36. Savage CR, Jutras BL, Bestor A, Tilly K, Rosa PA, Tourand Y, Stewart PE, Brissette CA, Stevenson B. 2018. Borrelia burgdorferi SpoVG DNA- and RNA-Binding Protein Modulates the Physiology of the Lyme Disease Spirochete. Journal of Bacteriology 200:e00033–18.

37. Barbour AG. 1984. Isolation and cultivation of Lyme disease spirochetes. Yale J Biol Med 57:521–525.

38. Yang XF, Pal U, Alani SM, Fikrig E, Norgard MV. 2004. Essential role for OspA/B in the life cycle of the Lyme disease spirochete. J Exp Med 199:641–648.

39. Livak KJ, Schmittgen TD. 2001. Analysis of relative gene expression data using real-time quantitative PCR and the 2-ΔΔCT Method. Methods 25:402–8.

40. Blevins JS, Revel AT, Smith AH, Bachlani GN, Norgard MV. 2007. Adaptation of a luciferase gene reporter and lac expression system to Borrelia burgdorferi. Appl Environ Microbiol 73:1501–1513.

41. Groshong AM, Gibbons NE, Yang XF, Blevins JS. 2012. Rrp2, a prokaryotic enhancer-like binding protein, is essential for viability of Borrelia burgdorferi. J Bacteriol 194:3336–42.

42. Bunikis I, Kutschan-Bunikis S, Bonde M, Bergström S. 2011. Multiplex PCR as a tool for validating plasmid content of Borrelia burgdorferi. J Microbiol Methods 86:243–247.

43. Hellman LM, Fried MG. 2007. Electrophoretic mobility shift assay (EMSA) for detecting protein-nucleic acid interactions. Nat Protocols 2:1849–1861.

44. He M, Zhang JJ, Ye M, Lou Y, Yang XF. 2014. Cyclic Di-GMP receptor PlzA controls virulence gene expression through RpoS in Borrelia burgdorferi. Infect Immun 82:445–52.

45. Groshong AM, Gibbons NE, Yang XF, Blevins JS. 2012. Rrp2, a prokaryotic enhancer-like binding protein, is essential for viability of Borrelia burgdorferi. J Bacteriol 194:3336–3342.

46. Ouyang Z, Blevins JS, Norgard MV. 2008. Transcriptional interplay among the regulators Rrp2, RpoN and RpoS in Borrelia burgdorferi. Microbiology 154:2641–2658.

47. Ouyang Z, Blevins JS, Norgard MV. 2008. Transcriptional interplay among the regulators Rrp2, RpoN, and RpoS in Borrelia burgdorferi. Microbiology 154:2641–2658.

